# Improving effect size estimation and statistical power with multi-echo fMRI and its impact on understanding the neural systems supporting mentalizing

**DOI:** 10.1101/017350

**Authors:** Michael V. Lombardo, Bonnie Auyeung, Rosemary J. Holt, Jack Waldman, Amber N. V. Ruigrok, Natasha Mooney, Edward T. Bullmore, Simon Baron-Cohen, Prantik Kundu

**Affiliations:** Department of Psychology, University of Cyprus, Cyprus; Center for Applied Neuroscience, University of Cyprus, Cyprus; Autism Research Centre, Department of Psychiatry, University of Cambridge, UK; Department of Psychology, School of Philosophy, Psychology, and Language Sciences, University of Edinburgh, UK; Brain Mapping Unit, Department of Psychiatry, University of Cambridge, UK; Section on Advanced Functional Neuroimaging, Departments of Radiology & Psychiatry, Icahn School of Medicine at Mount Sinai, USA

**Keywords:** multi-echo EPI, statistical power, denoising, task-fMRI, mentalizing, cerebellum

## Abstract

Functional magnetic resonance imaging (fMRI) research is routinely criticized for being statistically underpowered due to characteristically small sample sizes and much larger sample sizes are being increasingly recommended. Additionally, various sources of artifact inherent in fMRI data can have detrimental impact on effect size estimates and statistical power. Here we show how specific removal of non-BOLD artifacts can improve effect size estimation and statistical power in task-fMRI contexts, with particular application to the social-cognitive domain of mentalizing/theory of mind. Non-BOLD variability identification and removal is achieved in a biophysical and statistically principled manner by combining multi-echo fMRI acquisition and independent components analysis (ME-ICA). Group-level effect size estimates on two different mentalizing tasks were enhanced by ME-ICA at a median rate of 24% in regions canonically associated with mentalizing, while much more substantial boosts (40-149%) were observed in non-canonical cerebellar areas. This effect size boosting is primarily a consequence of reduction of non-BOLD noise at the subject-level, which then translates into consequent reductions in between-subject variance at the group-level. Power simulations demonstrate that enhanced effect size enables highly-powered studies at traditional sample sizes. Cerebellar effects observed after applying ME-ICA may be unobservable with conventional imaging at traditional sample sizes. Thus, ME-ICA allows for principled design-agnostic non-BOLD artifact removal that can substantially improve effect size estimates and statistical power in task-fMRI contexts. ME-ICA could help issues regarding statistical power and non-BOLD noise and enable potential for novel discovery of aspects of brain organization that are currently under-appreciated and not well understood.

---

A common criticism of neuroscience research in general (Button et al., 2013) and functional MRI (fMRI) in particular (Yarkoni, 2009), is that studies are characteristically statistically underpowered. Low statistical power by definition means that a study will have less of a chance for detecting true effects, but also means that observed statistically significant effects are less likely to be true and will be more susceptible to the biasing impact of questionable research practices (Button et al., 2013; Ioannidis, 2005). This problem is important given the emergent ‘crisis of confidence’ across many domains of science (e.g., psychology, neuroscience), stemming from low frequency of replication and the pervasive nature of questionable research practices (Button et al., 2013; Ioannidis, 2005; Simmons et al., 2011).

Low statistical power can be attributed to small sample sizes, small effect sizes, or a combination of both. The general recommended solution is primarily to increase sample size (though other secondary recommendations also include increased within-subject scan time). These recommendations are pragmatic mainly because these variables are within the control of the researcher during study design. While these recommendations are important to consider (Desmond and Glover, 2002; Friston, 2012; Lindquist et al., 2013; Mumford and Nichols, 2008; Yarkoni, 2009), other considerations such as dealing with substantial sources of non-BOLD noise inherent in fMRI data also need to be evaluated before the field assumes increasing sample size or scan time to be the primary or only means of increasing statistical power. These considerations are especially poignant when mandates for large-N studies and increased within-subject scan time are practically limiting due to often cited reasons such as the prohibitively high costs for all but the most well-funded research groups or in situations where the focus is on studying sensitive, rare, and/or less prevalent patient populations and where increasing scan time is impractical (e.g., children, neurological patients).

On the issue of non-BOLD noise variability, it is well known that fMRI data are of variable quality. Poor and variable quality data can significantly hamper ability to achieve accurate and reproducible representations of brain organization. It is widely understood that the poor sensitivity of fMRI often arises from high levels of subject motion (often task correlated), cardiopulmonary physiology, or other types of imaging artifact (Murphy et al., 2013). These artifacts are problematic because they are often inadequately separable from the functional blood oxygenation level dependent (BOLD) signal when using conventional fMRI methods. Given an advance in fMRI methodology that allows enhanced detection and removal of these artifacts, the situation regarding statistical power and sample size may change markedly. Such advances could create viable experimental alternatives or supplements to the recommendation for increasing sample size/scan time to boost statistical power, and concurrently make for an fMRI approach that can more reliably enable discovery of subtle but potentially key aspects of typical and atypical brain function.

In this study, we address problems related to statistical power through specific targeting of the problems related to non-BOLD artifact variability. We have applied an approach that integrates the fMRI data acquisition innovation of multi-echo EPI with the decomposition method of independent components analysis (ICA), towards principled removal of non-BOLD signals from fMRI data. Our fully integrated implementation is called multi-echo independent components analysis or ME-ICA (Kundu et al., 2012). ME-ICA utilizes multi-echo fMRI to acquire both fMRI signal time series *and their NMR signal decay*, towards distinguishing functional BOLD from non-BOLD signal components based on their respective and differentiable signatures in the decay domain. Critically, BOLD and non-BOLD signal domains are readily differentiable in data analysis of the echo time (TE) domain – irrespective of overlap of signal patterns in the spatial and temporal domains. BOLD-related signals specifically show linear dependence of percent signal change on TE, whereas non-BOLD signal amplitudes demonstrate TE-independence. Therefore, ME-ICA is a biophysically and statistically principled bottom-up approach towards identifying and retaining BOLD-related variability while systematically removing non-BOLD variation. ME-ICA has been successfully applied to fMRI resting state connectivity acquisition and analysis and is shown to enable various improvements with regards to increased temporal signal-to-noise ratio (tSNR), enhanced ability to remove motion and other artifactual sources of variability, more principled statistical modeling in seed-based connectivity analysis, enhanced specificity, and translational capacities for use at high-field strength and within animal models (Kundu et al., 2015; Kundu et al., 2013; Kundu et al., 2014). ME-ICA can also be applied alongside multi-band acquisition (Olafsson et al., 2015) and has recently been applied to identify ultra-slow temporally-extended task-related responses (Evans et al., 2015). However, one very necessary yet unexamined niche within the space of uses for fMRI is within the highly utilized context of traditional task-based fMRI studies and the potential impact that ME-ICA innovations could have on effect size estimation and consequently statistical power.

Here we conduct the first assessment of how ME-ICA performs with regards to effect size estimation and statistical power in task-related activation mapping settings with block-designs. ME-ICA can be flexibly applied to both task- and resting state fMRI contexts. This unified approach is advantageous since conventional resting state and task data processing and denoising use disjoint pipelines that often may require different technical skillsets. However, it is important and currently not well understood if and to what extent generalized non-BOLD removal as targeted by ME-ICA enhances the elucidation of task effects, compared to current task activation analysis with inline denoising based on linear artifact models and arbitrary filtering done in a study specific manner. Ultimately, because task-fMRI approaches form the bedrock of our understanding of human brain function and it is not clear how ME-ICA’s principled removal of non-BOLD signal could impact final inferences in such studies. In this study we specifically examined how ME-ICA performs against a conventional task-based imaging analysis pipeline with regressing out motion parameters (i.e. TSOC+MotReg) and another prominent yet more recent task-based denoising procedure (i.e GLMdenoise (Kay et al., 2013)). We utilized two separate tasks (i.e. the ‘SelfOther’ and ‘Stories’ tasks) tapping neural systems supporting the social cognitive domain or mentalizing and theory of mind and highlight its effects in terms of effect size estimation and statistical power. We also evaluate the impact of the method on two sets of brain regions; ‘canonical’ regions typically highlighted as important in the neural systems for mentalizing (Frith and Frith, 2003; Lombardo et al., 2010b; Saxe and Powell, 2006; Schaafsma et al., 2015; Schurz et al., 2014; Spunt and Adolphs, 2014; van Overwalle, 2009) and ‘non-canonical’ regions in the cerebellum (van Overwalle et al., 2014).

## Materials and Methods

### Participants

This study was approved by the Essex 1 National Research Ethics committee. Parents gave informed consent for their child to participate and each child also gave assent to participate. Participants were 69 adolescents (34 males, 35 females, mean age = 15.45 years, sd age = 0.99 years, range = 13.22-17.18 years) sampled from a larger cohort of individuals whose mothers underwent amniocentesis during pregnancy for clinical reasons (i.e. screening for chromosomal abnormalities). The main focus for sampling from this cohort was to study the fetal programming effects of steroid hormones on adolescent brain and behavioral development. At amniocentesis, none of the individuals screened positive for any chromosomal abnormalities and were thus considered typically developing. Upon recruitment for this particular study, we additionally checked for any self- or parent-reported neuropsychiatric conditions. One individual had a diagnosis on the autism spectrum. The remaining participants did not have any other kind of neurological or psychiatric diagnosis. Analyses were done on the full sample of 69 individuals, as analyses leaving out the one patient with an autism diagnosis did not change any of the results.

### Task Design

Participants were scanned using two block-design fMRI paradigms. The first paradigm, which we call the ‘SelfOther’ task, is a 2 x 2 within-subjects factorial design which contains two contrasts that tapped either self-referential cognition and mentalizing and was similar in nature to previously published studies (Lombardo et al., 2011; Lombardo et al., 2010a; Lombardo et al., 2010b). Briefly, participants are asked to make reflective judgments about either themselves or the British Queen that varied as either a mentalistic (e.g., “How likely are [you/the Queen] to think that it is important to keep a journal?”) or physical judgment (e.g., “How likely are [you/the Queen] to have bony elbows?”). Participants make judgments on a 1-4 scale, where 1 indicated ‘not at all likely’ and 4 indicated ‘very likely’. All stimuli are taken from Jason Mitchell’s lab and have been used in prior studies on mentalizing and self-referential cognition (Jenkins et al., 2008; Mitchell et al., 2006). The SelfOther task is presented in 2 scanning runs (8:42 duration per run; 261 volumes per run). Within each scanning run there are 4 blocks per condition, and within each block there are 4 trials of 4 seconds duration each. Task blocks are separated from each other by a 16 second fixation block. The first 5 volumes of each run are discarded to allow for equilibration effects.

The second paradigm, which we call the ‘Stories’ task, is also a block-design and contains two contrasts tapping mentalizing and language domains. The paradigm is identical to a study by Gweon and colleagues (Gweon et al., 2012), utilizing the same stimuli and presentation scripts provided directly by Gweon and colleagues. Briefly, participants listen to a series of stories whereby the stories differ in content. The content of the stories vary in terms of mentalistic, social, or physical content. The social stories contain descriptions of people and characters but make no statements that referenced mental states. Physical stories are segments of stories that describe the physical setting but do not include people. Mental stories are segments that include references to people as main characters and make references to mental states that those characters hold. The paradigm also includes blocks for two other kinds of language control conditions that are not examined in this manuscript (i.e. stories read in a foreign language (e.g., Russian, Hebrew, and Korean) and blocks of music played by different instruments (e.g., guitar, piano, saxophone, and violin)). After participants heard each story segment they were given a choice of whether a specific auditory segment logically came next. This was introduced to verify that participants were paying close attention to the stories and the details inside each story segment. The Stories task is presented in 2 scanning runs (6:36 duration per run; 192 volumes per run) and within each scanning run there are 2 blocks per condition. The first 6 volumes were discarded to allow for equilibration effects.

Resting state data was also collected on each participant with a 10 minute long ‘eyes-open’ run (i.e. 300 volumes), where participants were asked to stare at a central fixation cross and to not fall asleep. The multi-echo EPI sequence was identical to those used in the task paradigms.

### fMRI Data Acquisition

All MRI scanning took place on a 3T Siemens Tim Trio MRI scanner at the Wolfson Brain Imaging Centre in Cambridge, UK. Functional imaging data during task and rest was acquired with a multi-echo EPI sequence with online reconstruction (repetition time (TR), 2000 ms; field of view (FOV), 240 mm; 28 oblique slices, descending alternating slice acquisition, slice thickness 3.8 mm; TE = 13, 31, and 48 ms, GRAPPA acceleration factor 2, BW=2368 Hz/pixel, flip angle, 90°). Anatomical images were acquired using a T1-weighted magnetization prepared rapid gradient echo (MPRAGE) sequence for warping purposes (TR, 2300 ms; TI, 900 ms; TE, 2.98 ms; flip angle, 9°, matrix 256 × 256 × 256, field-of-view 25.6 cm).

### fMRI Preprocessing

Data were processed by ME-ICA using the tool *meica.py* as distributed in the AFNI neuroimaging suite (v2.5 beta10), which implemented both basic fMRI image preprocessing and decomposition-based denoising. For the processing of each subject, first the anatomical image was skull-stripped and then warped nonlinearly to the MNI anatomical template using AFNI *3dQWarp*. The warp field was saved for later application to functional data. For each functional dataset, the first TE dataset was used to compute parameters of motion correction and anatomical-functional coregistration, and the first volume after equilibration was used as the base EPI image. Matrices for de-obliquing and six-parameter rigid body motion correction were computed. Then, 12-parameter affine anatomical-functional coregistration was computed using the local Pearson correlation (LPC) cost function, using the gray matter segment of the EPI base image computed with AFNI *3dSeg* as the LPC weight mask. Matrices for de-obliquing, motion correction, and anatomical-functional coregistration were combined with the standard space non-linear warp field to create a single warp for functional data. The dataset of each TE was then slice-time corrected and spatially aligned through application of the alignment matrix, and the total nonlinear warp was applied to the dataset of each TE. No time series filtering was applied in the preprocessing phase. Data were analyzed both with no spatial smoothing and with a 6mm full-width-half-maximum (FWHM) spatial filter. The effective smoothness from second-level group analyses are reported for each task and each analysis in Supplementary Table 2.

Multi-echo fMRI data enables analysis where time series of different TEs and thus different BOLD contrasts can be combined to synthesize time series of optimal contrast specific to each voxel, a process called “optimal combination.” For a given voxel with a particular T2* value, the signal acquisition of optimal BOLD contrast is at TE=T2*. While conventional fMRI involves data acquisition at a single TE (selected to reflect the average tissue T2*), some voxels have T2*>TE and others have T2*>TE, meaning BOLD contrast-to-noise ratio is not homogeneous throughout the brain. The greatest macro-inhomogeneities are near areas of high magnetic susceptibility such as orbitofrontal cortex or temporal bone, where conventional fMRI suffers substantial signal “drop-out.” However, with multi-echo fMRI, each voxel’s T2* can be estimated specifically, and a given voxel’s time series of different TEs can be averaged with weights to produce a computed signal time series with BOLD contrast approximating an acquisition of TE=T2*, done for every voxel. This procedure first involves computing an empirical map of magnetic susceptibility, where for each voxel, the TE-specific signal time series means are fit to an exponential decay model, with the rate constant parameter being the T2* estimate. The voxel-specific T2* values are then used to calculate weights for averaging time series across different TEs, with each voxel- and TE-specific weight being calculated as (Posse et al., 1999):

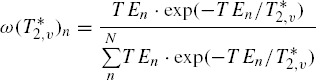

This procedure implemented a matched-filter that produced a contrast-optimized or “high dynamic range” time series dataset where the functional contrast-to-noise at each voxel was maximized and thermal noise is attenuated. The “optimally combined” time series dataset (abbreviated TSOC) was used in all further analysis steps (i.e. ME-ICA, TSOC+MotReg, GLMdenoise). Note that the TSOC dataset is input into all denoising procedures (i.e. after basic preprocessing) to ensure a fair comparison across techniques. While ME-ICA denoises TSOC data by further exploitation of multi-echo data through TE-dependence/independence analysis, other pipelines instead attempt to remove noise primarily through the inclusion of noise regressors in the first-level GLM (i.e. motion regressors or global noise regressors).

### ME-ICA Denoising

Time series denoising with ME-ICA was based on ICA decomposition of optimally combined multi-echo data and component classification informed by signal models reflecting the BOLD-like or artifact-like signal decay processes. This has been detailed in our prior work (Evans et al., 2015; Kundu et al., 2013) and is summarized in Supplementary Table 1. The decomposition path utilized in ME-ICA is designed to elucidate components specifically with the BOLD TE-dependence pattern, but is implemented similarly to other ICA treatments: dimensionality is first reduced using a PCA step, then spatial ICA is applied to dimensionally reduced data to find sparse or statistically independent sources (e.g. in MELODIC PICA, dimensionality reduction is achieved with probabilistic PCA, followed by FastICA). Dimensionality reduction is important for reducing the complexity of the ICA problem (to an optimal number of components depending on the data), and concomitantly reducing the proportion of Gaussian-distributed noise (explained by many low-eigenvalue components) that would otherwise cause the ICA solution to fail in convergence. However, ME-ICA differs from other ICA treatments in the dimensionality reduction step. Conventional automatic dimensionality estimation cannot analyze individual principal components for their mechanisms of signal origin, and instead utilizes assumptions on the statistical distribution of noise. In contrast, ME-ICA implements ME-PCA, a model-based approach to distinguishing principal components representing MR contrast versus thermal noise as, respectively, those components with high Kappa, Rho, or variance relative to corresponding spectrum elbows vs. those with none of these properties. While high-variance principal components are preserved for subsequent ICA decomposition by both conventional and ME-PCA, the latter retains low-variance components with MR contrast. In this way, ME-PCA achieves higher dimensional ICA decompositions, indicating potentially greater sensitivity in elucidating BOLD components in ICA decomposition, while utilizing a more direct detection of Gaussian distributed thermal noise, indicating potentially greater specificity and ICA stability for higher dimensional (i.e. more comprehensive) ICA solutions. Following dimensionality reduction based on ME-PCA, ME-ICA applies spatial FastICA using the *tanh* contrast function to identify a spatial basis of statistically independent component maps, alongside a complementary matrix of time courses (the mixing matrix).

The mixing matrix was fit to the time series of each separate TE, producing coefficient maps for each component at each TE. The signal scaling of each component across TEs was then used to compute Kappa (*κ*) and Rho (*ρ*), which were pseudo-F statistics indicating component-level TE-dependence and TE-independence, respectively. While it is understood that ICA separates data into statistically independent components, for multi-echo fMRI these metrics were evaluated to determine the segregation of signals into components of specifically BOLD-related or non-BOLD related contrasts (1 and 2, in Supplementary Table 1), indicating a higher order decomposition than the one achieved by ME-PCA which produced mixed-contrast components (5) versus thermal noise. In addition to BOLD and non-BOLD groups, a usually small group of mixed BOLD/non-BOLD components related to draining vein physiology (4) are elucidated and rejected as not neuronally related. Finally, for time series denoising, the full mixing matrix (including all component time courses) is fit to the optimally combined (i.e. the source data for the ME-PCA/ICA decomposition pipeline) with multiple linear least squares regression, and the time series fit corresponding to rejected components submodel is subtracted from the optimally combined time series. The number of components selected for all subjects on all runs of both tasks can be found in Supplementary Table 2.

### Task-fMRI Data Analysis

All first and second level statistical modeling was performed in SPM8 (http://www.fil.ion.ucl.ac.uk/spm/), using the general linear model (GLM). First level analyses modeled the hemodynamic response function (HRF) with the canonical HRF, and used a high-pass filter of 1/128 Hz. In contrast to ME-ICA, we also ran denoising with two other prominent approaches; GLMdenoise (Kay et al., 2013) and via conventional task-based fMRI analysis that included motion regressors in the first-level GLM model (TSOC+MotReg). It is important to re-iterate that each pipeline (ME-ICA, GLMdenoise, and TSOC+MotReg) utilized TSOC data. For GLMdenoise, global noise regressors are identified with cross validation across runs and used as regressors of no interest in first-level individual subject GLMs. For TSOC+MotReg we mimicked conventional task-based fMRI analysis by using motion parameters as regressors of no interest in first-level individual subject GLMs. When running first-level GLMs on ME-ICA denoised data, we did not include motion parameters as regressors of no interest because such artifact is already removed in principled manner at the prior denoising step. All first-level individual subject GLMs modeled the specific contrast of Mentalizing>Physical, and these contrast images were input into second-level random effects GLM analyses (i.e. one sample t-test). Any whole-brain second-level group analyses we report are thresholded at a voxel-wise FDR q<0.05 (Genovese et al., 2002).

### Resting State fMRI Connectivity Analysis

Resting state connectivity on ME-ICA processed data was estimated using the multi-echo independent components regression (ME-ICR) technique developed by Kundu and colleagues (Kundu et al., 2013). This analysis technique effectively controls for false positives in connectivity estimation by using the number of independent components estimated by ME-ICA as the effective degrees of freedom in single-subject connectivity estimation. Once ME-ICA has the estimated number of components, these component maps are concatenated, and connectivity is estimated by computing the correlation of ICA coefficients between the seed and other brain voxels. The seed regions we have chosen are the peak voxels from the NeuroSynth ‘mentalizing’ map in right and left hemisphere cerebellum (RH MNI x = 29, y = -82, z = -39; LH MNI x = -25, y = -78, z = -39). Connectivity GLM analyses were implemented within SPM and the second-level group connectivity maps are thresholded with a voxel-wise FDR threshold of q<0.05.

To assess the similarity between whole-brain resting state connectivity and Mentalizing>Physical task-activation maps, we used robust regression (Wager et al., 2005) to compute the correlation between the whole-brain connectivity and activation maps. Robust regression allows for protection against the effects of outliers that are particularly pronounced in the connectivity maps, since voxels that contain or are proximally close to the seed voxel exhibit very large connectivity values.

In contrast to connectivity estimated via ME-ICA data with ME-ICR, we also ran conventional functional connectivity analyses on the TSOC data. Here we followed standard analysis procedures such as bandpass filtering and motion regression. These steps are achieved using AFNI *3dBandpass* to bandpass filter the data between 0.01 and 0.1 Hz, after orthogonalizing data with respect to a baseline (motion parameters, etc.) matrix (*-ort* argument) to additionally remove motion-related variability all in one step. No other steps were taken to denoise the data (e.g., global signal regression, white matter regression, etc). The bandpass filtered and motion-regressed data were then inserted into GLMs in SPM8. Note here that bandpass filtering was only applied in this analysis of conventional resting state connectivity analysis and was not done in ME-ICA and ME-ICR connectivity analyses.

To compare the difference between activation-connectivity correlations for ME-ICR vs TSOC+MotReg, we use the *paired.r* function within the *psych* R library (http://cran.r-project.org/web/packages/psych/) to obtain z-statistics to describe the difference between correlations. However, no hypothesis tests (i.e. p-values) are computed for these analyses as they are not needed since the comparisons are on correlations estimated from the entire population of interest (i.e. all voxels within whole-brain volume).

### Effect Size Estimation and Power Simulations

All effect size and power estimates were computed with the *fmripower* MATLAB toolbox (http://fmripower.org) (Mumford and Nichols, 2008). Effect size is operationalized here as a standardized measure of distance from 0 expressed in standard deviation units (i.e. mean/sd) and is analogous to Cohen’s d. Here the mean refers to the contrast image (i.e. con*.img) produced by the second-level random effects analysis. The standard deviation is taken by computing the square root of the variance image (i.e. ResMS.img) produced by the second-level random effects analysis. We have made one change to the code within *fmripower* in how it computes effect size. This change allows us to compute effect size at each voxel and then to average the effect size across ROI voxels. This is different from the current implementation in *fmripower* which will first compute the average mean and standard deviation values across ROI voxels and then computes effect size based on these average mean and standard deviation values. Within *fmripower* the Type I error was set to 0.05 and we computed power across a sample size range from n=5 to n=100. All effect size and power estimates were estimated from independently defined meta-analytic ROIs identified by NeuroSynth (http://neurosynth.org) (Yarkoni et al., 2011) for the feature ‘mentalizing’. This feature contained 98 studies and 4526 activations. The NeuroSynth ‘mentalizing’ mask was first resampled to the same voxel sizes as the current fMRI datasets. Because regions surviving the NeuroSynth analysis at FDR q<0.01 were large and contained multiple peaks (e.g., medial prefrontal cortex comprised both dorsal and ventral subregions), we constrained ROIs further by finding peak voxels within each region, and constructing a 8mm sphere around each peak. This resulted in 11 separate ROIs. Eight of the 11 have been reported and heavily emphasized in the literature (dorsomedial prefrontal cortex (dMPFC): x = -2, y = 60, z = 22; ventromedial prefrontal cortex (vMPFC): x = -2, y = 48, z = -20; right temporo-parietal junction (RTPJ): x = 59, y = -55, z = 27; left temporo-parietal junction (LTPJ): x = -48, y = -55, z = 26; posterior cingulate cortex/precuneus (PCC): x = 2, y = -52, z = 42; right anterior temporal lobe (rATL): x = 48, y = -6, z = -20; left anterior temporal lobe (lATL): x = -52, y = 6, z = -35; left temporal pole (lTP): x = -40, y = 21, z = -24). The remaining 3 regions are located in the cerebellum (right hemisphere cerebellar region Crus II (rCereb): x = 29, y = 82, z = -39; medial cerebellar region IX (mCereb): x = 2, y = -52, z = -47; left hemisphere cerebellar region Crus II (lCereb): x = -25, y = -78, z = -39) and have been relatively overlooked in the literature, with some exceptions that also rely on meta-analytic inference (van Overwalle et al., 2014).

To get an indication of how big the effect size boost due to ME-ICA was, we computed a measure of effect size percentage increase operationalized as (ES_ME-ICA_ – ES_TSOC or GLMdenoise_/abs(ES_TSOC or GLMdenoise_) * 100. Bootstrapping (1000 resamples) was then used to re-run SPM second-level group analysis and fmripower computations in order to construct 95% confidence intervals around effect size and effect size percentage increase (i.e. ‘effect size boost’) estimates. The calculation of confidence intervals for the effect size boost metric allowed us to determine which brain regions showed robust ME-ICA related effect size boosts compared to either GLMdenoise or TSOC+MotReg pipelines. Any region that showed a lower bound 95% confidence interval estimate above 0 was considered a region whereby ME-ICA robustly improves effect size estimation over and above GLMdenoise or TSOC+MotReg pipelines. In addition to this strict set of criteria using confidence intervals, we also report descriptively the percentage of bootstrap resamples whereby ME-ICA provided a larger effect size than GLMdenoise and TSOC+MotReg.

To further describe the effects of ME-ICA over and above GLMdenoise TSOC+MotReg pipelines, we have computed the minimum sample size to achieve 80% power and the sample size and cost reduction due to using ME-ICA to achieve a study with 80% power, assuming a per subject scanning cost of $500. In cost savings computations, any regions that did not achieve requisite power before n=100 were excluded from such calculations.

## Results

### ME-ICA Denoising on the Raw Time Series

Before touching on quantitative comparisons of effect size and power due to ME-ICA, it is helpful to convey properties of the images and time series acquired with ME acquisition, as well as the effect of ME-ICA denoising directly on the time series. ME sequences capture the decay of EPI images and (time series) with increasing TE, shown in Fig 1A. For example, ME data show the signal evolution of susceptibility artifact (i.e. signal dropout) in areas such as ventromedial prefrontal cortex (vMPFC) – it is made clear from Fig 1A that signal dropout occurs at longer TEs, as affected regions have short T2* due to proximity to air-tissue boundaries. Additionally, gray/white signal contrast increases over longer TE due to T2* differences between these tissue types. The T2*-weighted optimal combination (TSOC) implements a matched-filter of TE images yielding a new image time series with optimized contrast (TE~T2*) and compensation of susceptibility artifact by weighting towards the early TE in areas with short T2*. In Fig 1B we present time series data from ventromedial prefrontal cortex (vMPFC), posterior cingulate cortex/precuneus (PCC), and right cerebellum in order to demonstrate the effect of optimal combination on the time series, and then the effect of removing non-BOLD noise using ME-ICA relative to modeled task blocks. It is particularly apparent that ME-ICA, without prior information on task structure, recovers task-based block fluctuations while much of the middle echo, TSOC, and non-BOLD isolated signals carrying complex artifacts including drifts, step changes, and spikes.

**Fig 1:**
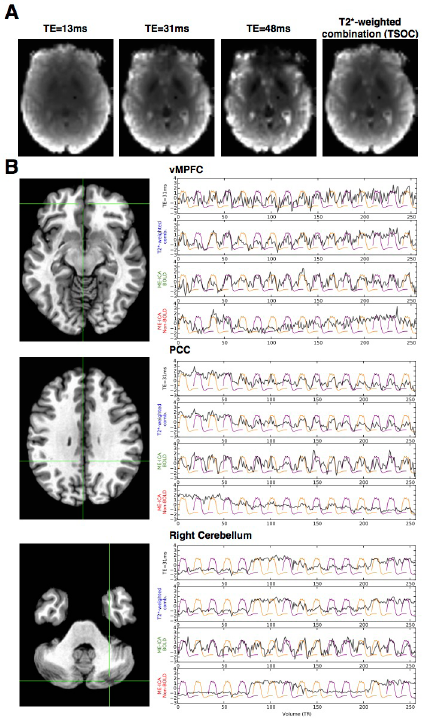
**Multi-Echo Signal Characterization.** Panel A shows the signal decay captured in multi-echo EPI images, for a single representative volume. With longer TE, gray/white contrast increases. Susceptibility artifact (e.g. dropout) also increases, as regions near in proximity to air-tissue boundaries have shorter T2*. The T2*-weighted optimal combination (TSOC) implements a matched-filter of TE images yielding a new image with optimized gray/white contrast (TE~T2*) and mitigation of susceptibility artifact. Panel B shows comparisons of time series data across three regions of interest; ventromedial prefrontal cortex (vMPFC), posterior cingulate cortex/precuneus (PCC), and right cerebellum. Each comparison shows the time series before model-based filtering of the middle TE image (black), TSOC image (blue), BOLD signals isolated on the basis of TE-dependence (green), and non-BOLD signals removed from the data (red). Purple and orange lines represent modeled mentalizing and physical blocks respectively.

### ME-ICA Boosts Effect Size Estimation

In evaluating ME-ICA-related effects on group-level inference, we examined the influence on non-BOLD denoising on effect size estimation. Effect size is operationalized here as a standardized measure of distance from 0 expressed in standard deviation units (i.e. mean/sd) and is analogous to Cohen’s d. As illustrated in Fig 2, with no smoothing (0mm) ME-ICA outperforms conventional methods for task-based fMRI analysis (TSOC+MotReg) and a prominent task-based denoising method (GLMdenoise) (Kay et al., 2013) (Fig 2B-C). This enhanced performance is evident across both mentalizing tasks and in nearly every single region investigated. Quantifying the magnitude of effect size boosting as the difference in effect size estimates, we find that the median ME-ICA induced boost for canonical mentalizing regions is around 24%. Boosts were much larger (nearly always greater than 50%) in areas such as vMPFC and left temporal pole (lTP) that characteristically suffer from signal dropout. Amongst cerebellar areas, right and left cerebellar Crus I/II areas showed evidence of even larger effect size boosts ranging from 48-149% increases when compared to GLMdenoise and 40-101% increases when compared to TSOC+MotReg. Under conditions where smoothing is done (i.e. 6mm FWHM) we see that many of the ME-ICA-related effect size boosts remain across several cerebellar and canonical cortical regions. However, this effect size boosting is smaller and less consistent across bootstrap resamples at 6mm FWHM, particularly within the SelfOther task. This phenomenon suggests that ME-ICA is relatively less dependent on smoothing in order to gain high degrees of sensitivity. In contrast, other methods likely require smoothing in order to enhance sensitivity. For example, it is clear that there are much larger increases in effect size at 6mm in GLMdenoise and TSOC+MotReg in the SelfOther task and such increases are not as prominent when going from unsmoothed data to 6mm data in ME-ICA. In other words, effect size estimates are more stable in ME-ICA in the SelfOther task, whereas in GLMdenoise and TSOC+MotReg, effect size estimates increase more (particular for cortical regions) simply due to smoothing. See Supplementary Tables 3-4 for full characterization of effect size estimates and effect size boosts.

**Fig 2:**
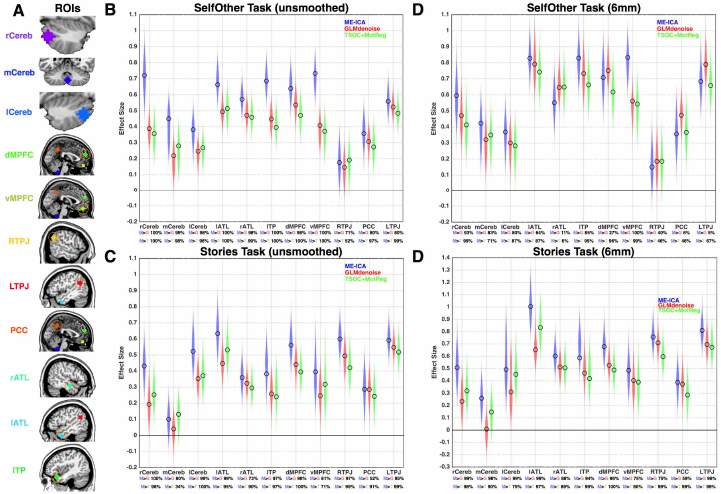
**ME-ICA Effect Size Boosting.** This figure shows effect size estimates from all regions of interest (panel A). Panels B and C show effect sizes in unsmoothed data, while panels D and E are from 6mm FWHM smoothed data Effect sizes are expressed in standard deviation units and are analogous to Cohen’s d. Colored clouds in each plot represent density of estimates obtained from 1000 bootstrap resamples, while unfilled black circles represent estimates within the true dataset. Below each region label on the x-axis are descriptive statistics indicating the percentage of bootstrap resamples where ME-ICA performed better than the alternatives (blue M, ME-ICA; red G, GLMdenoise; green T, TSOC+MotReg).

Because our operational definition of effect size is a standardized measure that incorporates both mean and variability measurements, we went further in decomposing how these boosts in effect size estimation manifested in terms of changes to either the mean and/or variability measurements. It is clear from Fig 3 that ME-ICA induces these boosts primarily by reducing estimates of variability at the 2^nd^ level group analysis. Given that at a within-subject level ME-ICA is working to remove non-BOLD noise from the time series, it is clear that one consequence of this for group-level modeling is clear reduction of between-subject variance which works to enhance standardized effect size estimates.

**Fig 3:**
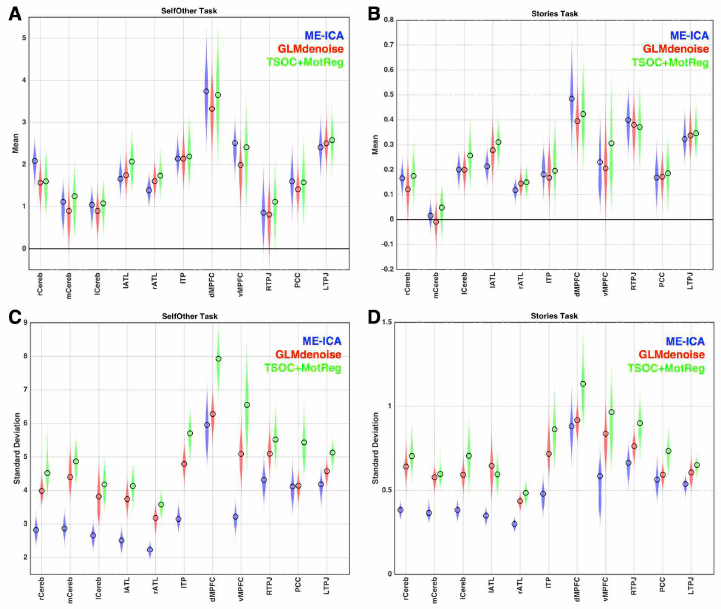
**ME-ICA Reduction in Variance in Group-Level Analyses.** This figure shows mean and standard deviation estimates from 2^nd^ level group modeling that contribute to the standardized effect size calculations. Panels A and B show mean estimates for all regions in both tasks. Panels C and D show standard deviation estimates. Colored clouds in each plot represent density of estimates obtained from 1000 bootstrap resamples, while unfilled circles represent estimates within the true dataset.

### Impact of ME-ICA on Statistical Power

Because ME-ICA improves standardized effect size estimation, it necessarily follows that statistical power will also be boosted, as such estimates are critical in such computations. However, for assessing the practical impact that ME-ICA may have, it is necessary to assess the impact such effect size boosting has on statistical power and sample size. Here we describe power simulations that mainly inform what we could expect in future work given effect size estimates similar to what we have observed in the current study under ME-ICA versus other analysis pipelines.

Power curves for each analysis pipeline across a range of sample sizes from n=5 to n=100 are illustrated in Fig 4A-B. Minimum sample size necessary for achieving 80% power at an alpha of 0.05 are shown in Fig 4C. Across all canonical regions and both tasks, the median minimum sample size to achieve 80% power at an alpha of 0.05 with ME-ICA is n=22. Minimum sample sizes across nearly all regions were well within reach of current standards for sample size (e.g., n<45). In contrast, for both GLMdenoise and TSOC+MotReg the median minimum sample size for canonical regions is n=38.

**Fig 4:**
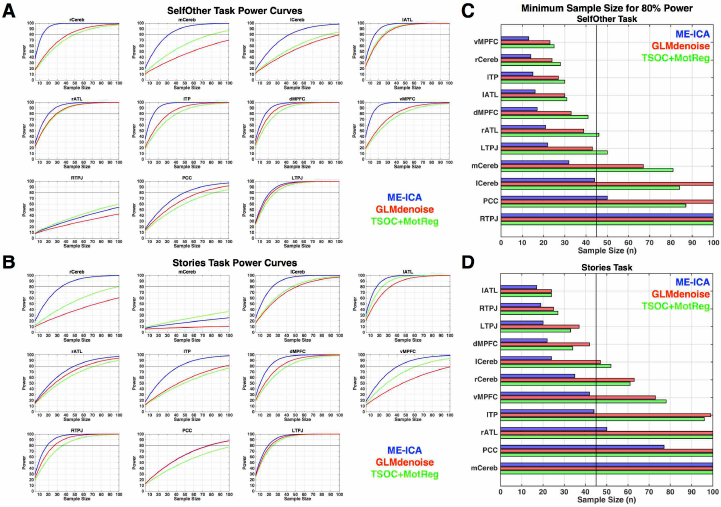
**Power Simulations**. This figure shows power curves constructed for each processing pipeline across a range of sample sizes from 5 to 100 (panels A-B). The minimum sample size necessary for achieving 80% power is shown in panel C for the Stories task (left) and SelfOther task (right). The dotted line indicates sample size of n=45.

For cerebellar regions, the power benefits due to ME-ICA were even more pronounced. Aside from medial cerebellar region XI (mCereb) in the Stories task which did not result in a sizeable effect (e.g., effect size <0.14), the minimum sample size needed for the bilateral cerebellar Crus I/II areas (rCereb, lCereb) were always well within the range of sample size that is typical for today’s standards when using ME-ICA (e.g., n<45). This stands in contrast to the situation for GLMdenoise and TSOC+MotReg, where sample size always required n>40 and in many instances was not attained by n=100.

For further illustration of practical impact, these boosts in statistical power and reduction in sample size necessary for achieving 80% power can be quantified into monetary savings. Assuming a scan rate of $500 per individual, if one was only interested in canonical regions, using ME-ICA would amount to median savings of $8,750 compared to GLMdenoise and $5,750 compared to TSOC+MotReg. If one was interested in cerebellar regions, using ME-ICA would amount to a median savings of $21,250 compared to GLMdenoise and $19,750 compared to TSOC+MotReg.

Visual examination of the power curves in Fig 4A-B highlights a point of diminishing returns when power is greater than 95%, as the improvements in power for adding more subjects diminishes substantially. We term this effect ‘saturation’. When using ME-ICA, many regions quickly reach these saturation levels at sample sizes that are practically attainable (e.g., n<45). In contrast, other pipelines like GLMdenoise and TSOC+MotReg typically require considerably larger sizes to hit these saturation levels.

### Functional Connectivity Evidence for Cerebellar Involvement in Neural Systems Supporting Mentalizing

The improvements in effect size estimation particularly for cerebellar regions is important as it potentially signals the ability of ME-ICA to uncover novel effects that may have been undetected in previous research. To further test the importance of cerebellar contributions to mentalizing, we have examined resting state functional connectivity data and the relationship that cerebellar connectivity patterns may have with task-evoked mentalizing systems. Prior work suggests that specific cerebellar regions may be integral to the default mode network (Buckner et al., 2011). The default mode network incorporates many of the regions that are highly characteristic in task-evoked systems supporting mentalizing (Andrews-Hanna et al., 2014). Meta-analytically defined cerebellar regions associated with mentalizing show some overlap with these cerebellar default mode areas (van Overwalle et al., 2015). Therefore, if cerebellar regions for which ME-ICA systematically produces boosts in effect size are integral in neural circuits associated with mentalizing, we hypothesized that resting state connectivity patterns with such cerebellar regions would be highly involved in the default mode network. Taking this hypothesis one step further, we also hypothesized that if these cerebellar nodes are truly important within the neural systems that support mentalizing, we should expect that cerebellar resting state functional connectivity patterns highlighted with multi-echo EPI methods would recapitulate the patterns observed for activational topology observed during mentalizing tasks across the whole-brain and within the same participants.

Confirming these hypotheses we find that bilateral cerebellar seeds involved in mentalizing show highly robust resting state functional connectivity patterns that resemble the default mode network within the same participants scanned on our task paradigms. Visually, the similarity between the ME-ICR connectivity maps and our Mentalizing>Physical activation maps are striking (Fig. 5A). Quantitatively we assessed this similarity through voxel-wise correlations (estimated with robust regression) across the whole-brain, and we confirm that the resting state functional connectivity maps are strikingly similar in patterning to what we observe for task-evoked mentalizing activation patterns (all r > 0.37) (Fig. 5B). Relative to the activation-connectivity similarity observed in TSOC+MotReg data, the activation-connectivity similarity obtained with ME-ICA and ME-ICR is much larger (i.e. z > 8.85) (Fig 5B-5D).

**Fig 5:**
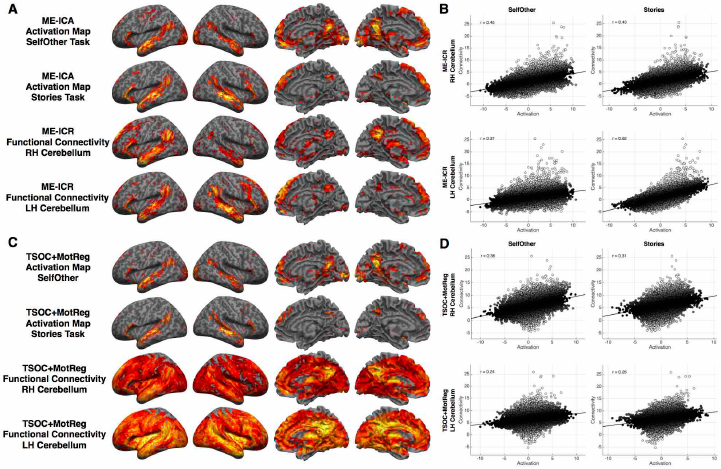
**Resting state functional connectivity from cerebellar seed regions and pattern similarity with Mentalizing>Physical activation maps.** This figure shows resting state connectivity from right and left cerebellar seed voxels (i.e. peak voxels from the NeuroSynth ‘mentalizing’ map) and their similarity to Mentalizing>Physical activation maps. Panel A shows activation and resting state functional connectivity maps when using ME-ICA and multi-echo independent components regression (ME-ICR). All data are visualized at thresholded of voxelwise FDR q<0.05. Panel B shows scatterplots and robust regression correlations between whole-brain activation and connectivity patterns when using ME-ICA and ME-ICR. Robust regression was used to calculate the correlation in a way that is insensitive to the outliers in the connectivity map which are voxels that are proximally close to the seed region. Panel C shows activation and cerebellar functional connectivity maps for data when using conventional analysis approaches on TSOC data. Activation maps are thresholded at FDR q<0.05. Connectivity maps are thresholded at the same t-statistic threshold for defining FDR q<0.05 in ME-ICR analyses (which were already much higher than the FDR q<0.05 cutoff estimated from TSOC data), and were shown in this manner to show connectivity at the exact same t-threshold cutoff. Panel D shows activation and connectivity similarity estimated with robust regression in TSOC data.

### Discussion

Task-based fMRI studies are characteristically of small sample size and thus likely underpowered for all but the largest and most robust effects. Furthermore, typical task-based fMRI studies do not apply advanced methods to mitigate substantial non-BOLD noise that is generally known to be inherent in such data. Combining small underpowered studies with little to no consideration of pervasive non-BOLD noise that is present in the data even after typical pre-processing and statistical modeling creates a situation where most task-based studies are potentially missing key effects and makes for somewhat impractical conditions for most researchers where massive sample sizes are required to overcome such limitations. In this study we show that ME-ICA results in robust increases in effect size estimation and statistical power in task-based block-design studies and these benefits tend to be most prominent without smoothing. These improvements are empirically demonstrated against other conventional and prominent analysis pipelines and denoising alternatives. As a consequence of these improvements in effect size and statistical power, we also demonstrate application of this method towards identification of novel effects in the cerebellum involved in the neural systems supporting mentalizing. Assuming similar effect sizes in future work, power simulations suggest that discovery of these novel cerebellar effects will remain nonetheless hidden at characteristically small sample sizes and without the multi-echo denoising innovations we report here.

ME-ICA-related benefits to effect size and statistical power are most prominent when data are unsmoothed. When utilizing standard smoothing kernels such as 6mm FWHM, boosts are still apparent but the gap between ME-ICA and other methods is smaller. This subtle effect related to smoothing may mean that other methods that do not benefit from ME-ICA denoising are less sensitive at higher spatial resolutions and will require some degree of smoothing in order to make up for that lack of sensitivity. On the other hand, one of the unique benefits with ME-ICA are the substantial increases in sensitivity even without smoothing and this may suggest that the powerful denoising ME-ICA implements could allow for much more sensitivity at finer-grained resolutions and in part may eliminate some need for smoothing. This advantageous characteristic may yield advantages in applying ME-ICA to other types of task-fMRI analysis such as multi-voxel pattern analysis (MVPA) (Norman et al., 2006) and representational similarity analysis (RSA) (Kriegeskorte et al., 2008), whereby added sensitivity at higher resolution is important and whereby the choice of smoothing is an important analytic consideration (Kamitani and Sawahata, 2010; Kriegeskorte et al., 2010; Op de Beeck, 2010). Given the potential for ME-ICA to enhance sensitivity without smoothing, it will be important in future work to explore whether ME-ICA to could enhance such applications of task-based fMRI analysis.

There are several practical points of impact that these results underscore. First, addressing the problem of statistical power in neuroscience, particularly fMRI studies (Button et al., 2013; Yarkoni, 2009), is a complicated matter as most recommendations for this problem rely on increasing the amount of data collected both at the within- and between-subject levels. A practical barrier for most research labs however, is that increasing the scale of data collection (e.g. massive sample size studies) is typically cost prohibitive. Our innovations here take a different perspective on the problem of low statistical power, by addressing from the bottom up, the problem of non-BOLD noise, which directly has impact on the sensitivity of fMRI, and thus statistical power. In practical terms, we show that ME-ICA allows for such substantial boosts in effect size estimation and consequently statistical power whereby in most cases (i.e. canonical and cerebellar regions investigated here), requisite levels of statistical power are attainable at sample sizes that should not be out of reach for most research laboratories. Therefore, if in the future researchers were to take up our multi-echo innovations in combination with uptake of already prominent considerations to generally collect more data, we could envision that the situation for fMRI research could substantially improve.

It is particularly important to underscore here that we are not suggesting that ME-ICA is the panacea to the small sample size problem and that as a result, researchers could continue the tradition of small sample size studies. Rather, we advocate that there are always compelling reasons to collect more data and that if funds permit, researchers should go above and beyond data collection that will ensure that their studies are highly powered at traditional sample sizes. Such a situation will ensure that canonical large effects are robust and replicable. Moreover, boosts in the sensitivity of fMRI can open up a range of previously practically unattainable possibilities for new discoveries. Such new discoveries could take the form of much more enhanced sensitivity for detecting smaller and more subtle effects in brain regions that are currently not well understood or which are methodologically hampered by being continually veiled underneath blankets of non-BOLD noise. New discoveries could also be enabled with parsing apart further variability such as subgroups that may have important translational implications (Lombardo et al., 2015), parsing apart heterogeneity mapped onto individual differences (Laumann et al., 2015), and/or more fine grained hypotheses/methods that result in much smaller effects than could be detected in the typical and more basic activation mapping paradigm. All of these situations could be substantially improved with a methodological approach that dramatically improves statistical power, but at the same time promotes and motivates researchers to collect larger samples than what is typically characteristic.

As an empirical demonstration of ME-ICA’s ability to enhance new discoveries for human brain functional organization, we have uncovered robust evidence that there are discrete cerebellar regions that should hold more prominence in discussions about the neural systems supporting mentalizing/theory of mind and the ‘social brain’. The cerebellum is already a neglected and not well-understood brain area, particularly in the context of its potential role in higher-level cognition (Buckner, 2013; Schmahmann, 1997; Stoodley and Schmahmann, 2009; van Overwalle et al., 2014; Wang et al., 2014). Prior indications that these cerebellar regions may be plausible candidates for neural systems supporting mentalizing come from meta-analytic evidence (van Overwalle et al., 2014). However, while meta-analytic evidence alone might suggest plausibility for these regions, it was still unclear as to the exact reasons for why these cerebellar regions have not been the topic of more extensive focus.

In this study, one of the novel findings that may help explain why these cerebellar regions are missed, is that they are typically veiled in substantial amounts of non-BOLD noise that obscure a researcher’s ability to detect such effects with traditional types of methods and analysis pipelines. Effect sizes for these regions under more traditional analysis approaches (e.g., TSOC+MotReg) are typically small and the sample size necessary for detecting those effects with high power are much greater than what is typical for fMRI research. However, after acquiring multi-echo data and applying ME-ICA, these effects are boosted by greater than 40%. As we show in this study, ME-ICA primarily boosts effect size estimation via noise reduction at the within-subject level and consequently has impact for reduction of variance at the group level. Therefore, it is clear that these regions are typically highly saturated in non-BOLD noise and this problem helps to obscure these effects from traditional research practices of using small sample sizes and conventional fMRI acquisition and denoising procedures that do not fully identify and remove such non-BOLD noise variability.

The ME-ICA application we present here should help researchers to gain a more stable foothold on cerebellar effects in the context of mentalizing and enable better circumstances for parsing apart how their role can further our understanding of such complex social cognitive processes. A promising avenue for future work on this topic would be to further understand the computational role the cerebellum plays in simulative processes that may be important in mentalizing (Ito, 2008; Mitchell et al., 2006). Translationally, the link between cerebellum and mentalizing is also particularly intriguing, given the longstanding, yet independent, literatures in autism regarding the cerebellum (Courchesne et al., 1988) and mentalizing (Baron-Cohen et al., 1985). Wang and colleagues have recently argued that atypical developmental processes within the cerebellum may be particularly important for understanding autism (Wang et al., 2014). Autism is well known for hallmark deficits in the domain of social-communication (Lai et al., 2014) and impairments in the development of mentalizing/theory of mind and self-referential cognition in autism (Lombardo and Baron-Cohen, 2010, 2011) as well as atypical functioning of neural mechanisms that bolster such processes (Lombardo et al., 2011; Lombardo et al., 2010a) are thought to be important as explanations behind social-communication deficits in autism. Thus, the intersection of developmental abnormalities in cerebellar development and their relationship to the development of mentalizing in autism will be an interesting new avenue of research enabled by these kinds of novel discoveries.

An important caveat for this study is that our findings are based on block-design activation paradigms, utilizing relatively long-duration changes in susceptibility weighting. This differs from event-related paradigms, whereby activations may be associated with a significant inflow component that is S0-weighted. Future studies will involve assessing the suitability of ME-ICA for the analysis of event-related studies as well as other more novel task-designs. With regard to novel task-designs such as temporally extended tasks, we have previously shown that ME-ICA also has the ability to separate ultra-slow BOLD effects from slow non-BOLD effects (Evans et al., 2015), and this opens up a range of possibilities for new paradigms that may be particularly well-suited for temporally-extended and continuous tasks, such as more naturalistic paradigms for social cognition (Schilbach et al., 2013; Zaki et al., 2009).

One limitation of the current study is the lack of comparison between multi-echo and traditional single-echo fMRI as well as emerging single-echo multi-band data. While our prior publications suggest that optimally combined multi-echo data is at least a fair proxy to single-echo data, there is some chance that the present comparison is conservative regarding the benefits of ME-ICA. We have previously shown that optimally combined time series data (TSOC) can readily double signal-to-noise ratio relative to unaccelerated single-echo fMRI (Kundu et al., 2013), via homogenizing functional contrast across the brain while attenuating thermal noise (combination is a weighted average implementing a matched- filter). Thus, in our view the most conservative comparison to make against ME-ICA should also be multi-echo data that benefits from enhanced tSNR over and above single-echo data. Recent work by Kirilina and colleagues provides direct comparisons of single- and multi-echo data in a task-fMRI context. These authors found that both kinds of acquisitions produced very similar group-level results in a task-fMRI context (Kirilina et al., 2016). This suggests that there is little benefit in group-level analyses for multi-echo acquisition (without ME-ICA) over and above traditional single-echo acquisitions. Applied to our work, these observations would suggest that analyses on TSOC data (i.e. TSOC+MotReg, GLMdenoise) are a useful approximation of what would be expected if we also compared ME-ICA directly to single-echo data. However, given all these important caveats, it will be important for future work on this topic to directly make this comparison of ME-ICA to single-echo data in task-fMRI contexts to confirm this prediction. Our prediction would be that given there is a boost in tSNR simply by acquiring multi-echo data and utilizing our T2* optimal combination method (Kundu et al., 2013) that ME-ICA would similarly outperform single-echo data.

The multi-echo innovations we provide here offer substantial improvements that can largely affect how the field conducts fMRI research. All of the tools for implementing these innovations are open source and most contemporary imaging facilities possess all the requisite requirements to enable actively taking up these innovations as standard practice. We hope that the community will actively take up these new innovations, as they are likely to have massive benefits for improving major issues that hamper the field and may further enable potential for new discoveries about human brain function.

## Acknowledgments

This work was supported by a Wellcome Trust project grant to SB-C and ETB. MVL was supported by the Wellcome Trust and fellowships from Jesus College, Cambridge and the British Academy. PK was supported by the National Institutes of Health–Cambridge Scholars Program. ETB is employed half-time by the University of Cambridge and half-time by GlaxoSmithKline (GSK).

## Legends for Supplementary Figures and Tables

**Supplementary Table 1: Classes of signal sources decomposed by ME-ICA.** Signal sources elucidated through combination of multivariate decomposition (PCA, ICA in order) and T2* decay analysis of multi-echo fMRI data as implemented in ME-ICA. κ is pseudo-F statistic component-level TE-dependent scaling suggesting network BOLD origin. ρ is pseudo-F statistic component-level TE-independent scaling suggesting artifact.

**Supplementary Table 2:** Number of BOLD-related components identified by ME-ICA for all subjects across both tasks. Effective smoothness of 2^nd^-level group analyses for each task, each analysis, and under 0mm or 6mm.

**Supplementary Table 3: Effect size, power, and effect size boosting statistics for data with no smoothing.** This table provides effect sizes and 95% CIs for all analysis pipelines and all regions when data is not smoothed. It also provides estimates of sample size needed to achieve 80% power. This table also provides information about effect size boosting statistics and 95% confidence intervals, impact on sample size estimates to achieve 80% power and impact on monetary savings.

**Supplementary Table 4: Effect size, power, and effect size boosting statistics for data with 6mm FWHM smoothing.** This table provides effect sizes and 95% CIs for all analysis pipelines and all regions when data is smoothed at 6mm FWHM. This table also provides information about effect size boosting statistics and 95% confidence intervals for all regions in both tasks.

